# Convergent genetic evolution within *Burkholderia multivorans* chronic infections correlates with lung function decline in cystic fibrosis patients

**DOI:** 10.64898/2026.09.16.751994

**Authors:** Mirela R. Ferreira, Sara C. Gomes, James E. A. Zlosnik, Vaughn S. Cooper, Leonilde M. Moreira

**Author notes:** Corresponding author: (VSC), (LMM).

## Abstract

The patterns and pathways of pathogen adaptation during chronic infections can reveal the gateways microbes must unlock to colonize and persist. In chronic lung infections of cystic fibrosis (CF) patients, a longtime goal has been to anticipate disease outcome and to improve treatment. Understanding of these processes for *Burkholderia multivorans,* the most prevalent CF pathogen of the *Burkholderia cepacia* complex, is limited. We investigate the evolution of seven different strains recovered from chronic airway infections of eight CF patients over 7-17 years. Despite divergent origins, each infecting population followed similar phylogenetic patterns of early diversification followed by the emergence of a dominant clade. The defining mutations of these prevalent lineages also affected the same set of global regulators, which altered clinically significant bacterial phenotypes in parallel directions. Mutated genes govern lipid metabolism, immune evasion, antibiotic resistance, biofilm production, and survival under oxygen limitation, and unite studies of chronic infections by different species of the *B. cepacia* complex. Most significantly, the phylodynamic signal that a dominant lineage had emerged within the infection, more so than any particular set of mutations, was associated with a more rapid decline in patient lung function. These findings reveal the importance of combining data from pathogen and host to link bacterial adaptation to disease progression, and ultimately to identify paths to reinforce host defense.

**Author summary:** Cystic fibrosis (CF) remains a serious disease despite major advances in care, because chronic lung infections can still gradually damage the airways and reduce lung function. Among the bacteria that can cause these long-term infections, the *Burkholderia cepacia* complex is especially concerning because it can survive many antibiotics and is associated with poor outcomes. In this study, we focused on *Burkholderia multivorans*, now one of the most common *Burkholderia* species found in CF lungs. By following bacterial populations from eight patients over several years, we found that these infections often evolved in a similar way: they first became genetically diverse, and later one dominant lineage emerged and took over. This transition was linked to changes in genes affecting antibiotic resistance, immune evasion, biofilm formation, and survival in low-oxygen conditions. Most importantly, the appearance of these dominant bacterial lineages was associated with faster decay of patient lung function. These findings show that bacterial evolution during chronic infections can follow predictable pathways that drive disease progression, which can enable earlier detection, monitoring, and treatment of CF lung infections.

## Introduction

The landscape of respiratory microbiology for cystic fibrosis (CF) patients has improved dramatically over the past few decades. Not only are patients living three times as long on average, the incidence and prevalence of airway infections that threaten well-being have declined steadily (1,2). One group of pathogens that is less common but which remains threatening is the *Burkholderia cepacia* complex (*Bc*c), which are 20+ closely related species that can establish long-term infections and survive most antibiotic treatments. The prevalence of *Bc*c infections has fallen from 3% to 1.2% in the US, and is now below 10% across Europe (3–6). Concurrently, the species distribution of *Bc*c infections has shifted globally towards *B. multivorans* and away from the typically more pathogenic *B. cenocepacia* (7). However, these changes resulted more from improved infection control preventing the sharing of transmissible strains and the advent of highly effective modulator therapy (i.e., elexacaftor/tezacaftor/ivacaftor) than from more focused pathogen diagnostics or treatment (6). As evidence, prevalence of *Bc*c among patients diagnosed with advanced lung disease is often 2-3 times as high (7). There remains a large unmet need to understand how opportunistic pathogens like *Bc*c that plague CF patients establish infections, persist throughout treatment, and ultimately threaten patient health.

The CF lung environment presents a range of novel stresses to bacteria, including dysregulated host immunity, frequent antibiotic exposure, altered nutrients, variable oxygen levels, and competition with other microorganisms (8). To colonize this complex environment and establish a chronic infection, a bacterial population must adapt to it by changes in gene expression and by mutations that improve response to the environmental demands (9–11). Some of these bacterial adaptations may enable persistence in novel niches within the CF airway that support long-term colonization and evade treatments, and some may ultimately cause disease and degrade lung function. These distinctions remain unclear partly owing to the challenge of gathering synchronized microbial samples and patient data over infections that may last decades. Studies of patient cohorts over time who are infected by similar pathogens hold great promise for improved understanding of the evolutionary pathways leading to infection chronicity, for better diagnostics, and for more optimal treatment.

Our understanding of pathogenic adaptation, or so-called “pathoadaptation”, has been mostly shaped by studies comparing the genomes and phenotypes of the predominant CF pathogen, *Pseudomonas aeruginosa* (8,12). Despite diverse mutated genes, genome sequencing of longitudinal isolates from CF patients consistently identified convergent mutations in antibiotic resistance, biofilm lifestyle, DNA repair, and global regulatory systems (8,9,13). Although these mutations in so-called pathoadaptive genes were found in separate infections caused by distinct strains, none of them appears to be solely responsible for predisposing *P. aeruginosa* towards chronic infections, suggesting multiple or combinatorial pathways to adaptation to the CF airway (14).

The notion that *P. aeruginosa* populations convergently evolved in the CF airway predates genome sequencing, as new bacterial phenotypes arose repeatedly in different patients (15,16). These traits tended to diversify early during chronic infection, followed by a consolidated set of pathoadapted characters after 2-3 years, including slower growth, reduced motility, aggregate formation, and elevated antibiotic resistance (8). More recent *in vivo* studies also show common shifts in metabolism, stress response, and nutrient scavenging across genetically diverse infections (17,18). Investigators and clinicians have thus sought to define bacterial traits that could serve as chronicity markers to guide distinct interventions (8).

By comparison, our understanding of chronic infections caused by *Bc*c in the CF airway is more limited. Patients are 10-20 fold less likely to be colonized, allowing fewer studies, and *Bc*c phenotypes are more variable and less definitive, partly because *Bc*c represents a cluster of ∼30 closely related species (19). Nonetheless, *Bc*c infections often contribute to morbidity and mortality and have been a criterion to exclude CF patients from life-saving interventions such as lung transplantation (20). One pioneering study of an outbreak of *B. dolosa* identified genes with parallel adaptive evolution because of strong, common selective pressures experienced within the CF airway (21,22), while a study of *B. cenocepacia* from multiple epidemic lineages revealed more diverse evolved genes and phenotypes, with some convergent phenotypic trajectories across patients (23).

Another *Bc*c species causing infections of the CF airway is *B. multivorans*, the focus of this study. Currently, *B. multivorans* is the most prevalent species infecting patients with CF in several countries (3–5,24). Although considered less virulent than *B. cenocepacia*, progressive decline in patient health associated with chronic infection by this bacterium is not uncommon (4,5,25). The first genomic study of a longitudinal *B. multivorans* series from a CF patient revealed initial diversification followed by emergence of a dominant clade that coincided with the decline in patient lung function and reduced airway microbial diversity (26). These isolates acquired multiple independent mutations in lipid metabolism genes, providing unequivocal evidence of strong selection. Later isolates also lost lipopolysaccharide O-antigen, motility, grew slower, while increasing antibiotic resistance, likely adaptations to chronicity (26). Two subsequent studies of *B. multivorans* evolving within the CF lung identified parallel genetic and phenotypic adaptations overlapping with prior studies (27,28), including pathways for antibiotic resistance and altered envelope biogenesis and metabolism. Nevertheless, an integrated approach linking genotypes, phenotypes, and clinical outcomes in a large-scale study remains an unmet goal.

To build a systematic understanding of how *B. multivorans* adaptively evolves when establishing and persisting in chronic infections of CF patients, we studied longitudinal changes among isolates from eight patients infected by diverse strains for periods between 7.1 and 17.6 years. We applied the latest genomic methods to characterize genetic changes within these infections and measured an array of relevant phenotypes to enable a genome-wide association study. Further, a combined analysis of bacterial genotypes and phenotypes with patient lung function data showed recurrent genotypic and phenotypic changes linked to a set of mutated genes predicted to enhance adaptation and persistence in the CF lung. Most notably, these analyzes suggest that worsening lung function was repeatedly associated with the emergence of a dominant clade of bacterial genotypes within the infecting population that adapted by multiple convergent traits.

## Results

### All but one patient were colonized by a single strain of *B. multivorans*

We studied 106 *B. multivorans* isolates from the sputum of 8 patients sampled between 1989 and 2015 by the Canadian *Burkholderia cepacia* Complex Research and Referral Repository (CBCCRRR) (Fig 1A and S1 Table). These samples were recovered over 7.1 to 17.6 years and include 6 to 21 isolates per patient. Patient attributes and co-infections detected are reported in Table 1. Changes in colony morphotypes, such as the mucoid-to-nonmucoid (or less mucoid) transition (29), motivated additional sampling. As shown in Fig 1A, some later isolates displayed reduced mucoidy. For most time points, only a single isolate was available, although for a few others, 2-4 independent isolates with visually distinct colony morphologies were maintained. These isolates were typed by RAPD and MLST assays, confirming that six of the patients were colonized by isolates belonging to a single strain, whereas patients P0431 and P0686 were initially colonized by two distinct strains with only one detected later. Patient P0686 was colonized by two distinct species, *B. multivorans* (2 isolates) and *B. pseudomultivorans* (9 isolates), which were not studied further. Patients P0280 and P0339 were colonized by isolates of the same RAPD/ST but all others were colonized by different strains (Fig 1A and S2 Table). We compared completely sequenced genomes of initial patient isolates with each other and with reference *B. multivorans* genomes. Four strains were closest to *B. multivorans* ATCC 17616, whereas the others were closer to ATCC BAA-247 (Fig 1B, S1 Fig and S3 Table), representing the two major *B. multivorans* clades (30). Strains within each clade differed by 1-2% in average nucleotide identity (ANI) and by 3% when comparing between clades (Fig 1B). Mobile elements were largely strain-specific, although some plasmids and prophages were shared across strains (S4 Table). Unique genes (BLASTP <75% similarity) with putative functions were linked to toxin-antitoxin systems, aromatic compound degradation, transporters, extracellular enzymes, and LPS O-antigen biosynthesis (S5 Table). In summary, each patient became colonized by 1-2 unique strains separated by considerable genomic distance, with one strain ultimately persisting and evolving to become less mucoid to varying levels.

**Fig 1.**
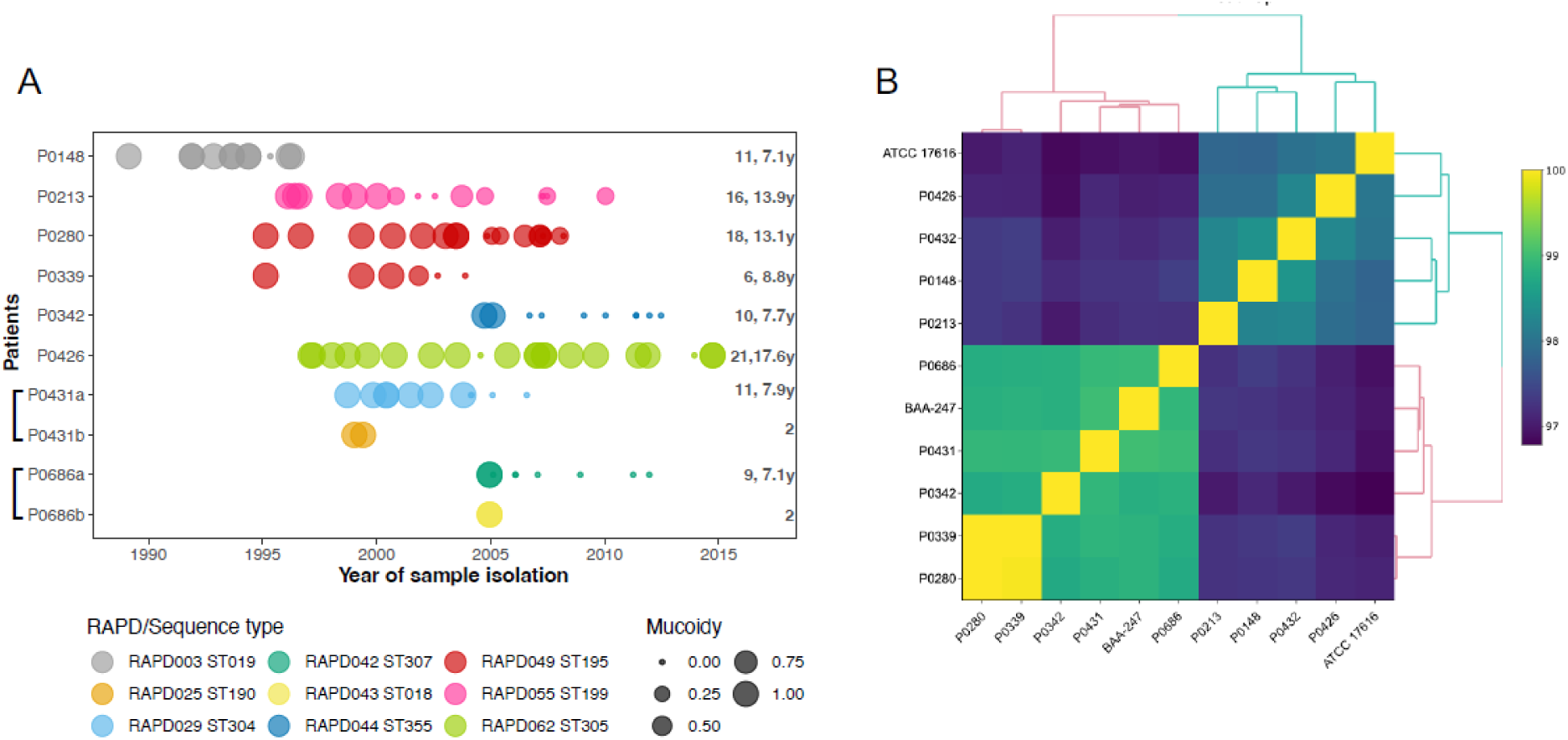
Longitudinal *B. multivorans* isolates collected from infections of eight CF patients over time. (A) Each patient was colonized by a different strain, defined by RAPD and Sequence Type (ST), except for patients P0431 and P0686 with two co-infecting strains (labeled “a” or “b”) early in their infections. Symbol size indicates level of colony mucoidy: nonmucoid (0), partially mucoid (0.25), mucoid (0.5-0.75), and frankly mucoid (1). The number of isolates and number of years (y) of sampling are shown at right for each infection. See S1 Table for more details. (B) Pairwise average nucleotide identity (ANI) among *B. multivorans* reference genomes ATCC 17616 and BAA-247 and the genome assemblies of the first isolates from each patient.

**Table 1.** Clinical descriptors of the patients under study and their frequencies of co-infections detected by culture-based methods.

| Parameter | Cases |
| --- | --- |
| no. | 8 |
| Median age at acquisition (range) | 13 (7-22) |
| Male, no. (%) | 3 (37.5) |
| CFTR genotype, no. (%) |  |
| ΔF508 homozygous | 6 (75) |
| Unknown | 2 (25) |
| Median number *B. multivorans* isolates (range) | 12 (6-21) |
| **Co-infections\*** | **no. (%)** |
| *Pseudomonas aeruginosa* | 7 (87.5) |
| *Staphylococcus aureus* | 8 (100) |
| *Haemophilus influenzae* | 6 (75) |
| *Aspergillus spp* | 5 (62.5) |
| *Enterobacter spp* | 4 (50) |
| *Candida spp.* | 3 (37.5) |
| *Stenotrophomonas maltophilia* | 2 (25) |
| *Serratia spp.* | 1 (12.5) |
| *Klebsiella spp* | 1 (12.5) |
| *Acinetobacter spp* | 1 (12.5) |
| *Alcaligenes spp* | 1 (12.5) |
| *Neisseria spp* | 1 (12.5) |
| *Mycobacterium spp* | 1 (12.5) |

\* at least one of these species was cultured and tested for antimicrobial susceptibilities during the period of their *B. multivorans* infection.

### Evolutionary diversification and lineage coexistence within patients inferred from clonal genomes

In theory, disparate colonizing strains could evolve along unique pathways when establishing chronic infection due to their founding genetic distinctions. Alternatively, strong selective pressures common to the CF airway could favor parallel genotypes and phenotypes across patients. To investigate these alternatives, we identified all newly arising mutations within each infection by comparison with the genome of the first isolate. This analysis revealed between 99 and 1392 genetic variants within patient series of ≥6 isolates, where P0342 exhibited the highest number of variant positions and P0148 the lowest (Table 2). SNPs and indels (<2 kbp) are shown in S6 Table, and large deletions (>2 kbp) in S7 Table. Mutation number depended on isolate count, infection duration, and mutations in DNA repair genes. Evolutionary rates (SNPs/year) varied considerably among infections: P0148 and P0426 isolates accumulated <4 SNPs/year, whereas those from P0280 and P0339 acquired 11.9 and 13.5 SNPs/year, respectively (S2A Fig), consistent with reported rates for the *Bc*c (2.1-15 SNPs/year) (21,26,27). Evolutionary rates in P0213 were variable due to a hypermutator isolate and a later, slower-evolving group, while P0342 and P0431a isolates showed higher rates driven by prevalent hypermutator lineages (S2A Fig and S6 Table). Excluding these hypermutators, the accumulation of SNPs over time within patients followed a linear trajectory (S2B Fig).

**Table 2.**
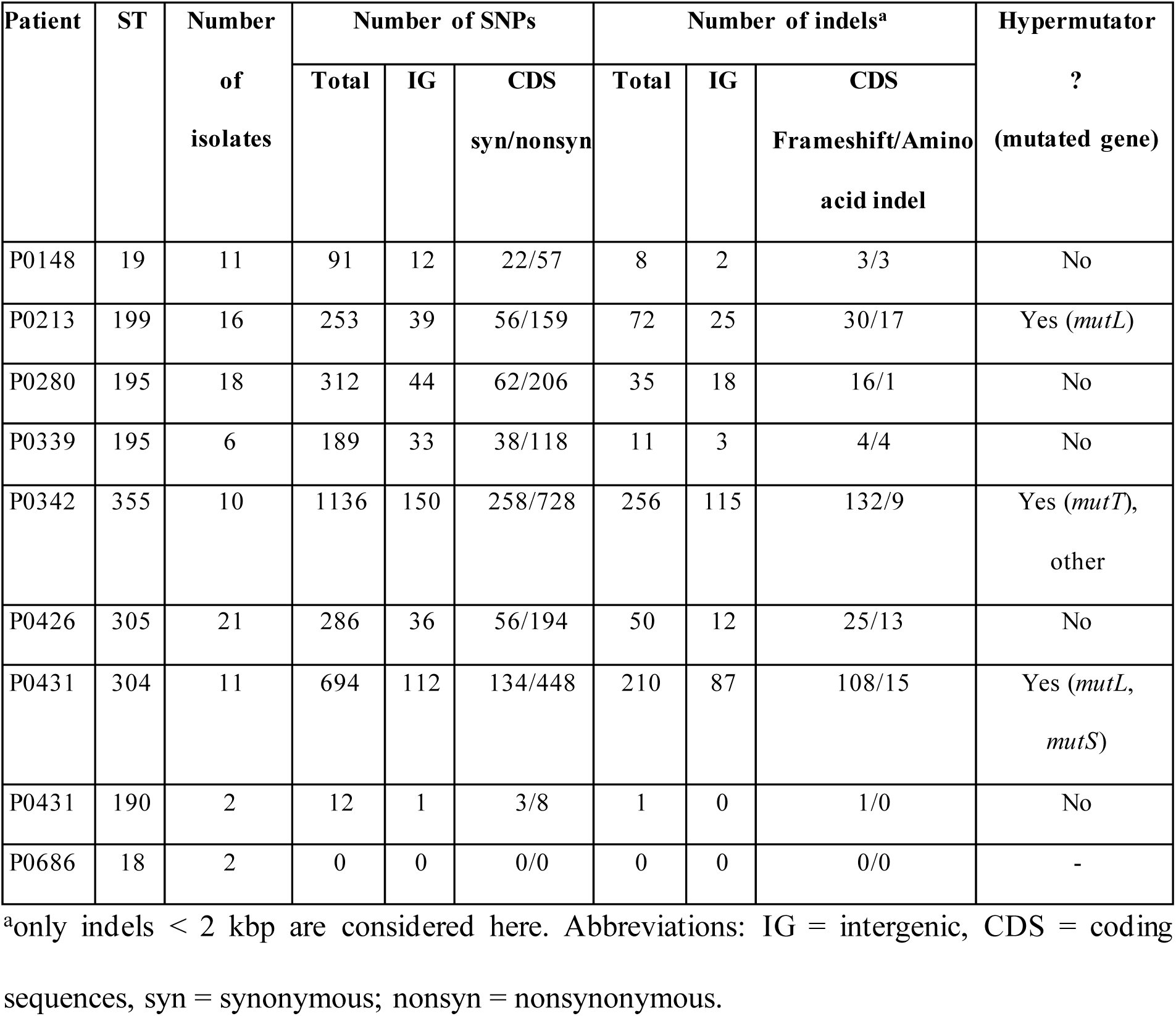
Number of mutations (SNPs and indels) identified within isolates from chronic *B. multivorans* infections.

Studies of chronic infections of the CF airway commonly show genetic and functional diversification due to spatial segregation and distinct ecology. Detailed phylogenies rather than overall evolutionary rates are required to capture these dynamics. We inferred within-patient phylogenies using the best-performing model including SNPs, indels, and large deletions. Two series with extensive sampling and lung function data covering the collection period are shown in Figs 2 and 3, with others in S3A-B Fig. The evolutionary dynamics within each patient appear to follow a remarkably consistent pattern: early isolates formed a tight cluster (clade C1), followed by the emergence of a dominant lineage (e.g., C3 in Fig 2; C2 in Fig 3) that displaced initial variants and diversified into sub-clades. Importantly, later isolates consistently belonged to dominant clades rather than earlier minor lineages (Figs 2 and 3; S3A-B Fig). Clade structure was assessed by comparing observed counts to a uniform expectation (chi-square) and by Kaplan-Meier persistence analyses. P0213 and P0280 series showed strong dominance (C3, *P*<0.0104; C2, *P*<0.0021, respectively), P0426 moderate dominance (C6, *P*<0.0092), and P0342 near monoclonality with 80% of isolates belonging to C2 (*P*<0.0558). Smaller series (≤6 isolates) lacked statistical power (S8 Table). Overall, despite genomic diversity and varying mutation rates, chronic *B. multivorans* infections shared consistent phylogenetic dynamics (phylodynamics) patterns.

**Fig 2.**
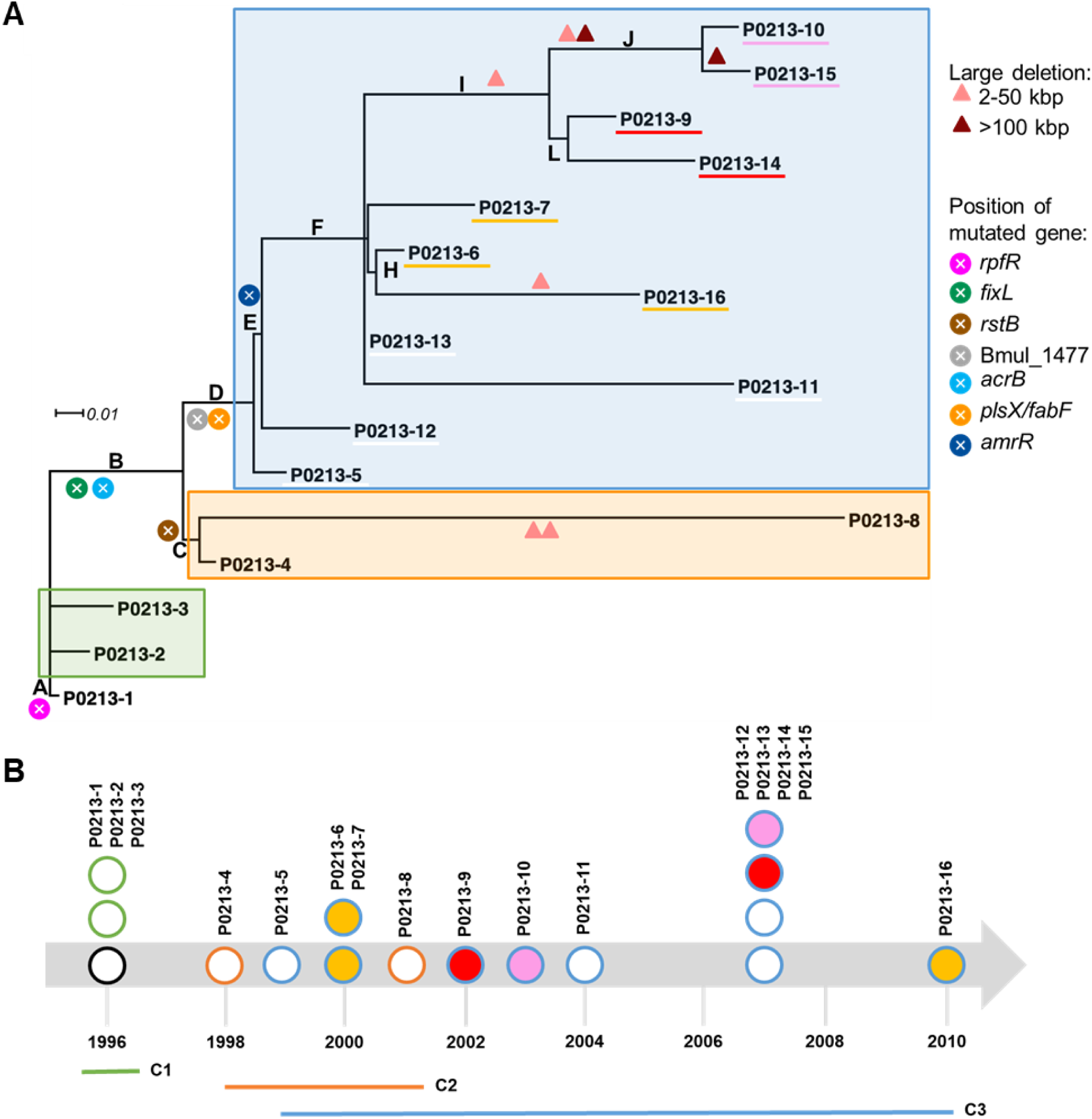
Genomic phylogeny reveals the emergence of a dominant clade that displaced earlier lineages within the infection. (A) Phylogeny of P0213 isolates modeling SNPs, indels and large structural variations in the genomes. The main branches are represented by capital letters, as defined in S10 Table. Symbols indicate the phylogenetic position of putative driver mutations occurring in the genes shown at right. (B) Temporal distribution of isolate sampling and their corresponding clades (C1, C2, C3) shows that clades coexisted. Symbol color indicates clade membership, with line color indicating major clade and fill color indicating minor clade, which is also denoted in (A) with colored underlines of isolate labels (clade C3 only).

**Fig 3.**
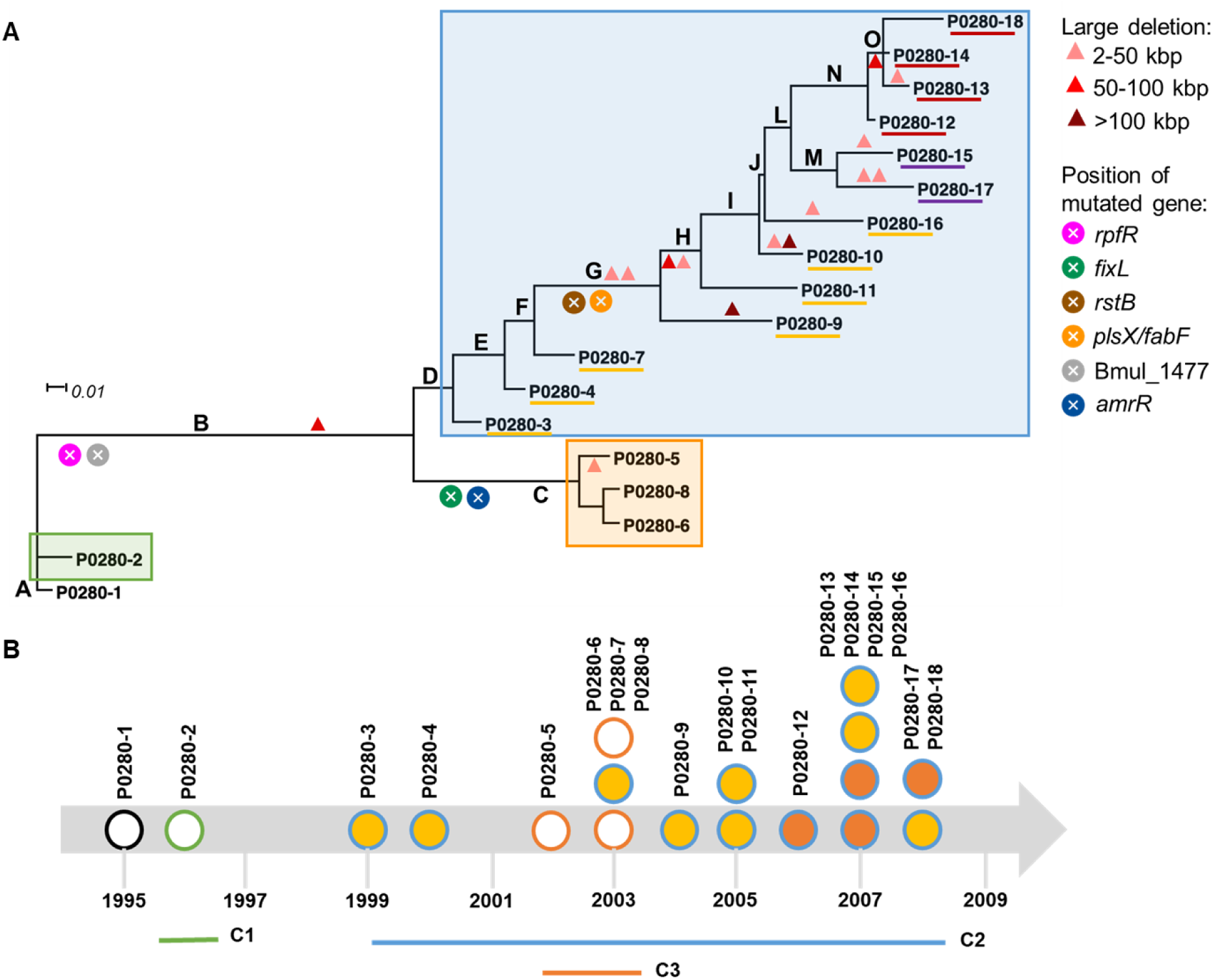
Genomic phylogeny reveals coexistence of diverse clades within a second patient and the rise of a dominant clade as the infection progresses. (A) Phylogenetic tree of P0280 isolates modeling SNPs, indels and large structural variations in the genome. The main branches are represented by a capital letter, as defined in S10 Table. Symbols indicate the phylogenetic position of mutations occurring in the genes shown at right. (B) Temporal distribution of isolate sampling and respective clades (C1, C2, C3) shows that clades coexisted. Symbol color indicates clade membership, with external line indicating major clade and fill color indicating minor clade, also shown as underlines of isolate labels in (A) (clade C2 only).

### Early diversification within infections is predominantly caused by single nucleotide changes but dominant lineages accumulate major deletions

Most early genetic diversification within each infection arose as SNPs, but the emergence of the dominant clades associated with large deletions (Figs 2-3, S3A-B Fig (colored triangles) and S6-7 and S9 Tables). Deletions were most prominent in P0280 and P0426 series, affecting secondary chromosomes (*B. multivorans* has three replicons) or a conserved plasmid and causing loss of multiple genes (S7 and S9 Tables). Most deletions involved hypothetical proteins of phage/plasmid origin (S4A-C Fig), including prophage regions in P0148 isolates 4, 5, 9 and in P0426, where one prophage was deleted in six isolates (S4A Fig). Nonetheless, some deletions affected virulence-related functions, including the major regulator of cyclic-di-GMP, *rpfR*, that responds to the signal *cis*-2-dodecenoic acid (in isolate P0426-21) and the *cepIR* genes encoding a quorum sensing system (in isolate P0213-8). A cluster of genes required for type VI secretion was lost from a clade within P0213 series, and several genes implicated in LPS O-antigen biosynthesis were lost from lineages within patients P0280 and P0426.

The emergence of hypermutator lineages is frequently reported in CF infections (26,31–33) and is thought to be an indirect response to strong selection on pathogens in fragmented populations where genetic variation is limiting (12). We identified mutator genotypes in 3 of 8 infections involving mutations in *mutS* and *mutL* genes in P0213 and P0431, respectively, while mutations in the putative DNA repair helicase PolX encoding gene and *mutT* were found in P0342 (Table 2 and S6 Table). However, hypermutators became dominant only in P0342 and P0431a. Taken together, small nucleotide changes such as SNPs and indels were most frequent during the early colonization period, while large deletions and the rise of hypermutators are more likely to occur as the dominant lineage became established and perhaps more spatially fragmented.

### Overwhelming evidence of parallel evolution across different patients

Because different *B. multivorans* strains exhibited similar evolutionary dynamics when infecting separate patients, we hypothesized that the CF airway exerted common selective pressures on certain bacterial traits that would produce shared genetic signatures. To test this, we assessed gene-level parallelism across longitudinal series, identifying genes mutated in >2 patients while excluding those with the same MLST/ST (P0280, P0339). In total, 108 genes showed parallel mutations (S10 Table), supporting shared host selection. This was strengthened by 55 genes with ≥4 independent mutations (subset in Fig 4). We evaluated the probability of this parallelism among non-hypermutator lineages using gene sizes, lineage-specific growth rates, and the genomic mutation rate (see Methods), and found the 29 genes in Fig 4 that were mutated in at least two patients remained statistically significant after correcting for multiple comparisons (FDR *q*<0.05; S10J Table). Among these, *plsX* was observed in all five series despite an expected hit frequency far below one per series, and *amrR*, Bmul_1477, *fabF*, *mpl*, and *relA* were repeatedly mutated in three or four series with *q*-values well below 0.05. These findings indicate that the recurrent genetic signatures are unlikely to reflect random mutations and instead support shared selective pressures in the CF airway.

**Fig 4.**
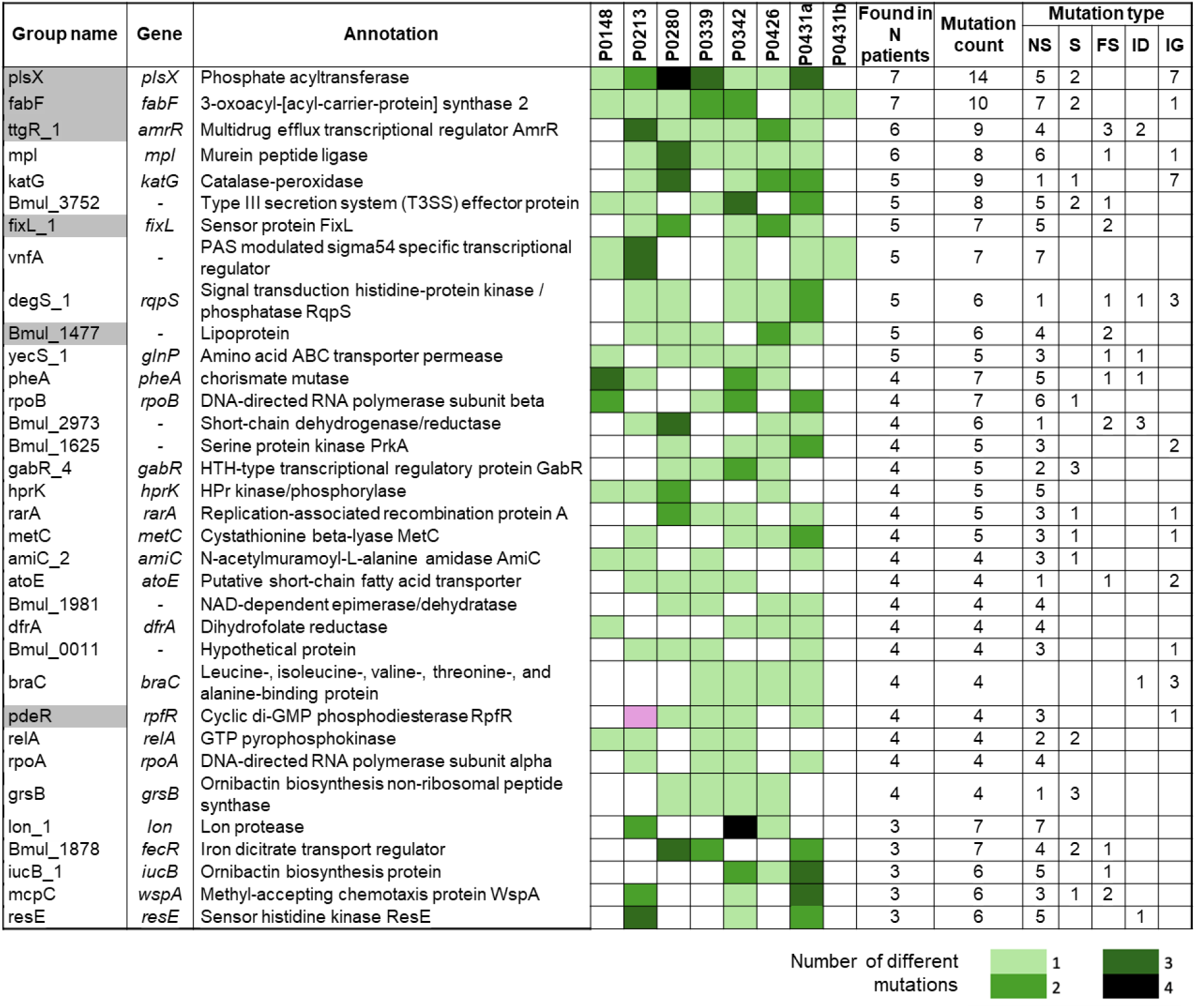
Exceptional convergent evolution of genes reveals common selection during chronic infection of the CF airway. Only genes with 3 or more independent mutational events are shown, irrespective of the total number of subsequent isolates with that mutation. Genes are annotated by ortholog cluster (group name) and a representative gene name is assigned when possible. The green color gradient indicates the number of different mutations within each patient. Mutation types: nonsynonymous (NS), synonymous (S), frameshift (FS), indels (ID), and intergenic (IG). Genes highlighted in gray accumulated mutations mostly at the phylogenetic base of early clades and are associated with lineage dominance. The pink box represents a nonsense mutation in the *rpfR* gene present in all isolates from P0213 series.

One of the two genes that became mutated in 7 out of 8 infections was *plsX* encoding phosphate acyltransferase, an enzyme involved in glycerophospholipid biosynthesis. The other gene was *fabF* which encodes 3-oxoacyl-[acyl-carrier-protein] synthase 2 required for fatty acids biosynthesis. Both belong to the same operon and are found under selection in other *Bc*c infections (22,23,26). Two additional genes became mutated in six patients: *amrR* encoding a TetR transcriptional repressor of an operon involved in aminoglycoside resistance (34), and *mpl* that contributes to peptidoglycan recycling and antibiotic resistance (35) (Fig 4). Seven more genes were mutated in five infections, including *fixL* encoding a sensor histidine kinase, previously linked to oxygen sensing and microaerophilic adaptation in *B. dolosa* and *B. multivorans* (21,26,36). Other recurrent targets involved oxidative stress, quorum sensing, gene expression regulation, transport, and antibiotic resistance (Fig 4 and S10 Table). Overall, most mutations were nonsynonymous or frameshift, likely impairing protein function, suggesting recurrent negative selection with occasional adaptive outcomes. To provide one example, mutations in the negative regulator of drug efflux, *amrR,* diminish function of this repressor protein and activate drug efflux constitutively (37).

### Mutated global regulators enable the evolution of dominant lineages during each infection

Populations arising from a single colonizing clone, as in most chronic CF infections (3,15,23,26), are shaped primarily by founder mutations shared by subsequent isolates rather than by changes confined to single isolates or minor lineages. To identify these potential driver mutations, we mapped all mutations to nodes or branches of the patient-specific phylogenies (Figs 2 and 3, S10A-G Table). Mutations that defined clades, or successful lineages, in most patients tended to be nonsynonymous. Across all mutations detected, 74-81% were nonsynonymous, exceeding the 72% expected under neutrality based on *Burkholderia* codon usage and G+C content (38). In contrast, clade-defining mutations shared by multiple isolates were 76-100% nonsynonymous, indicating stronger selection at these branch-defining events (S10H Table).

To identify putative driver mutations enabling lineage persistence during infection, we focused on genes mutated at the phylogenetic root of at least two dominant lineages. This identified 26 genes (Fig 4, S10I Table in gray) mainly linked to transcription/signal transduction and cell wall/membrane biogenesis. Six genes (Fig 4, gray) were mutated in dominant lineages in ≥4 patient series (see also Figs 2 and 3 and S3A-B Fig, colored dots represent specific genes). Notably, *rpfR* was frequently mutated, including in series P0280, P0339, P0342, and P0431a, and all P0213 isolates carried a nonsense mutation. This highlights RpfR as a global regulator integrating BDSF (cis-2-dodecenoic acid) signaling, cyclic-di-GMP, and key virulence/persistence traits (39,40). Global regulator *fixL* accumulated mutations in emergent lineages across multiple patient series (P0213, P0342, P0426, P0431a, P0280) while *rstB*, a sensor kinase encoding gene, is associated with dominant clades in P0280 and P0426 (Figs 2 and 3 and S3A-B Fig). This analysis also reinforces the role of *plsX*/*fabF* (lipid biosynthesis) and *amrR* (multidrug efflux repression), as most emergent lineages carried mutations in these genes. Together, this retrospective phylogenomics identifies global regulators (*rpfR*, *fixL, rstB*), lipid biosynthetic enzymes (*plsX*, *fabF*), and the efflux pump repressor *amrR* as key drivers of successful *B. multivorans* lineages in CF infections.

### Suite of progressive phenotypic changes occurred within nearly all infections

We conducted a broad array of phenotyping on all isolates to understand how *B. multivorans* traits evolved during long-term colonization of CF airways. These phenotypes were: mucoidy, LPS O-antigen presence, adhesion to CF lung epithelial cells, virulence against *Galleria mellonella* larvae, growth rate in synthetic CF medium (SCFM), susceptibility to antibiotics (aztreonam, piperacillin+tazobactam, kanamycin, and ciprofloxacin), biofilm formation, and swimming and swarming motilities (S1 and S11A Tables). These phenotypes were chosen because they tend to change during the course of the infection and are thought to enhance immune evasion, antimicrobial resistance and/or increase persistence under stress, and they have been shown to associate with declining patient lung function (23,26). Extensive variation in these phenotypes as well as temporal trends during the seven chronic infections are shown in Fig 5 and S5 Fig. Notably, progressive decreases in mucoidy, motility, antibiotics susceptibility, virulence to *Galleria*, growth in SCFM, and O-antigen presence were observed. Adhesion to CF epithelial cells generally increased (4 series increased and 1 decreased), and biofilm formation was variable (1 series increased, 3 decreased, and 3 did not show a tendency) and neither trend was statistically significant (*P*-value >0.1) (Fig 5). It is remarkable that this large-scale survey of 12 distinct phenotypes from nearly 100 isolates of different *B. multivorans* strains showed changes in parallel directions within six of seven patients. Taken together with the clear parallel genetic evolution, these phenotypes likely represent adaptations or correlates of adaptation to the CF environment.

**Fig 5.**
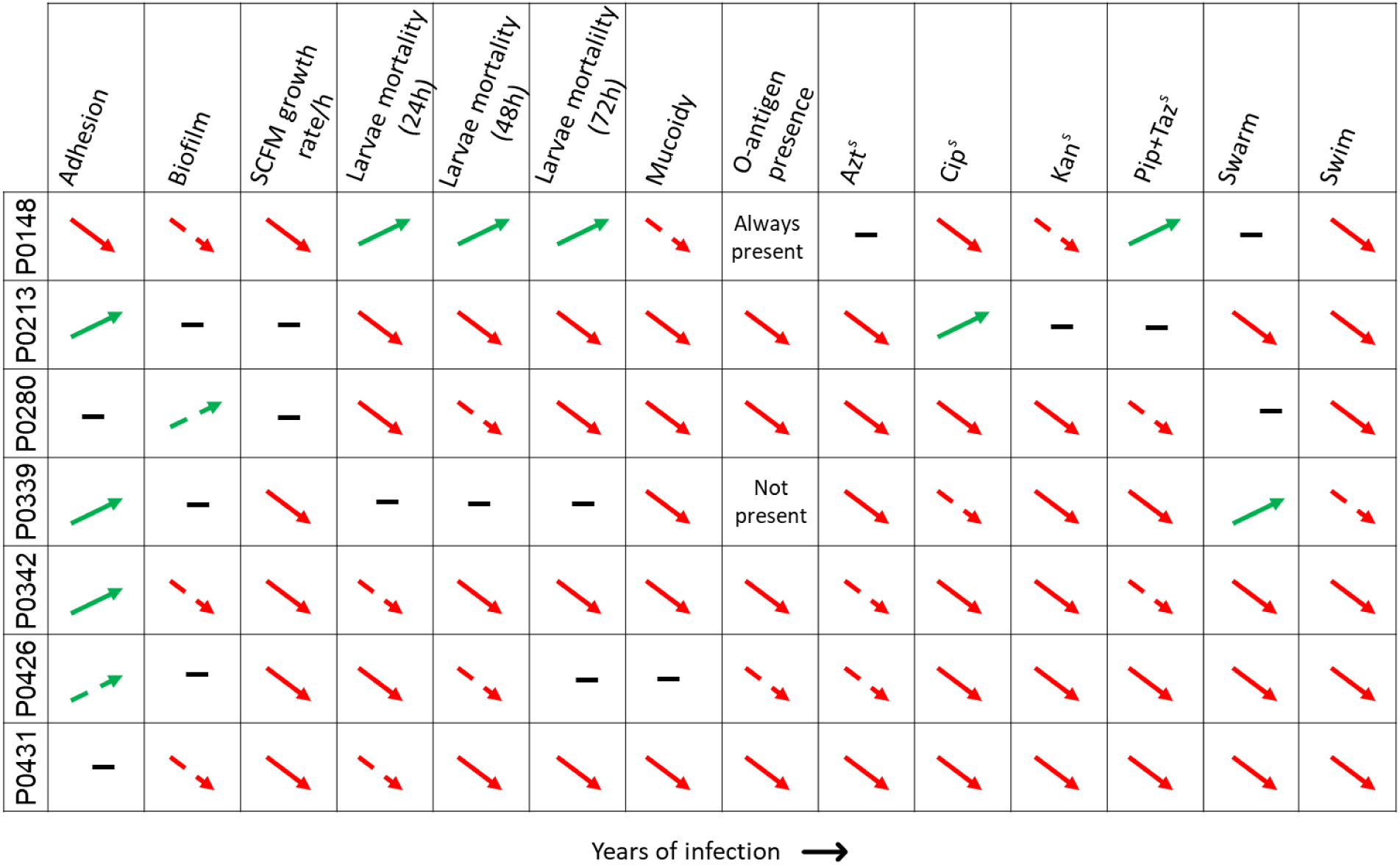
Clinically relevant phenotypes undergo progressive temporal changes during chronic infections. Spearman rank correlation was used to determine whether the phenotypes of isolates sampled over time from chronic infections changed over time. Solid arrows represent *p*-value <0.1; dashed arrows represent *P*-value >0.1; and (**–**) for no tendency. Azt^s^, susceptibility to aztreonam; Cip^s^, susceptibility to ciprofloxacin; Kan^s^, susceptibility to kanamycin; and Pip+Taz^s^, susceptibility to piperacillin+tazobactam.

### Genotype-phenotype association identifies new genes associated with *Burkholderia* pathoadaptation

To evaluate associations between prevalent genotypes (Fig 4) and changing bacterial phenotypes (Fig 5), we conducted a systematic genotype-phenotype association study (S11 and S12 Tables, see Methods for more detail). For each phenotype, data were simplified into two matrices: a binary genotype matrix defining the allele of each gene as “wild-type” or “mutant”, and a normalized quantitative matrix of phenotypic measurements. This analysis revealed several genetic predictors of clinically significant phenotypes (summarized in S13 Table with statistical evidence), beginning with the well-studied mucoid phenotype that declined in most patients. These mutated genes include *fabF* and *resE* (Fig 6A). FabF handles fatty acid elongation acting on bacterial membranes (41) and may also affect quorum sensing signals required for polysaccharide biosynthesis, whereas the uncharacterized ResE sensor kinase may regulate EPS production. Genes *bceC* and *wzc* affect synthesis of exopolysaccharide cepacian (42), validating our statistical approach (S13 Table).

**Fig 6.**
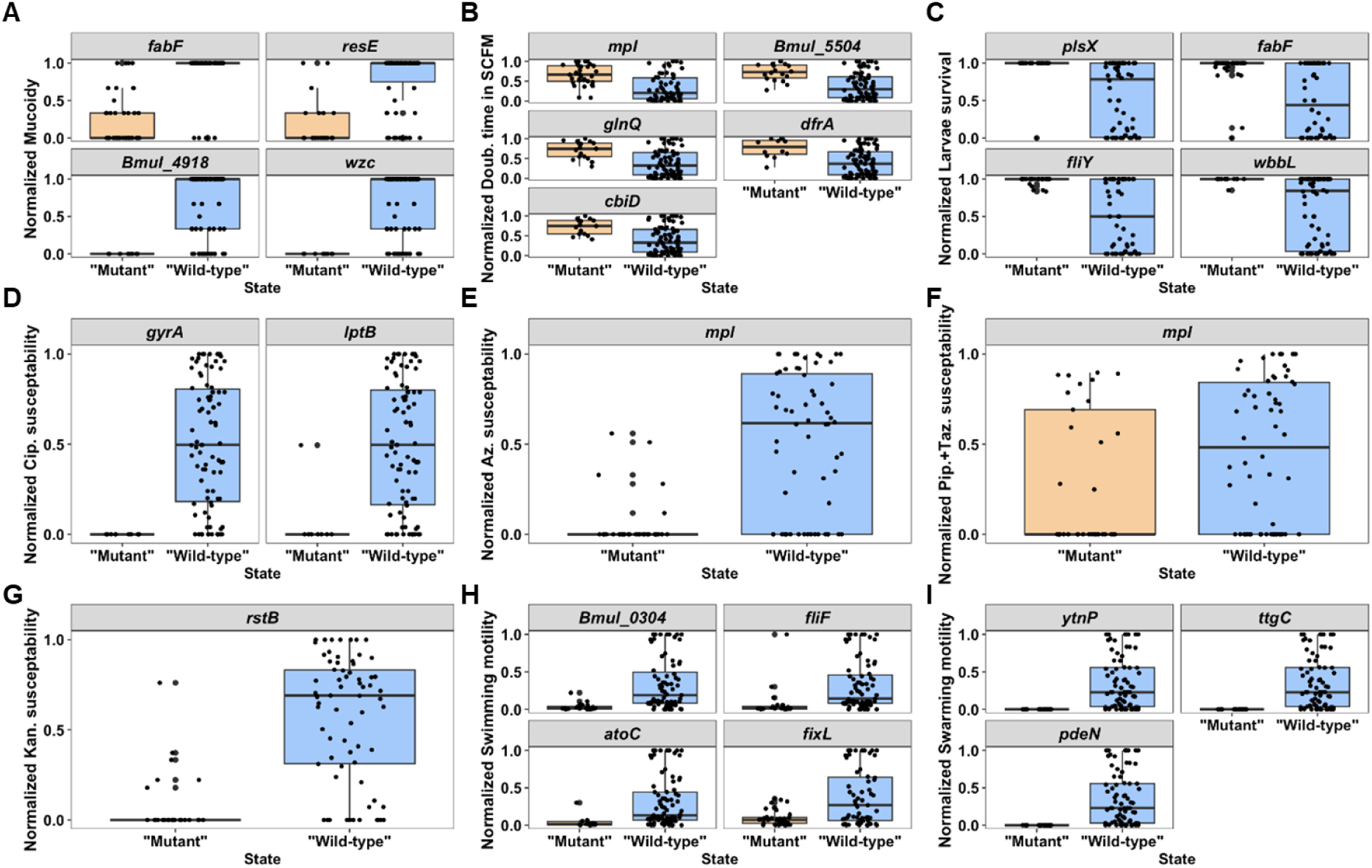
Genotype-phenotype association identifies genes linked to clinically relevant phenotypes. Systematic genotype-phenotype association analyses identifies several genes correlated with (A) mucoidy; (B) doubling time in SCFM; (C) acute virulence (larvae survival); (D) ciprofloxacin susceptibility; (E) aztreonam susceptibility; (F) piperacillin+tazobactam susceptibility; (G) kanamycin susceptibility; (H) swimming motility; and (I) swarming motility.

Genes associated with slower growth rates included *mpl* (Fig 6B) involved in peptidoglycan recycling, genes involved in trehalose metabolism (Bmul_5504), glutamine transport (*glnQ*), and cofactor biosynthesis (*dfrA* and *cbiD*). Genes associated with reduced virulence in *Galleria* are *plsX, fabF*, and *fliY* encoding a putative solute-binding protein (Fig 6C). In addition, mutations in *wbbL*, involved in LPS O-antigen biosynthesis, were associated with reduced virulence.

As expected, resistance to ciprofloxacin was correlated with mutations in *gyrA* but also to a putative transporter encoding gene *lptB* (Fig 6D). Mutations in *mpl* were associated with increased resistance to aztreonam and piperacillin-tazobactam, and *rstB* mutations were associated with kanamycin resistance (Fig 6E-G). While *gyrA* and *mpl* have established roles in resistance in several bacteria, including *Burkholderia* (21) and *Strenotrophomonas* (35), the contribution of these other identified genes remains unknown.

Loss of swimming motility was correlated with mutations in genes (e.g. *fliF*, *fixL*) well known to influence motility (Fig 6H), while reduced swarming motility was associated with mutations in genes that have not yet been characterized in *Burkholderia* (Fig 6I).

Spearman rank values of these associations and asssociated *P*-values are shown in S13 Table. Gene cluster naming follows Roary pangenome categories (S16 Table). Dots represent individual isolates that were categorized as mutant or wild-type depending on the presence or absence of a mutation in the indicated gene. The y-axis represents the value of the normalized phenotype for each isolate determined within each series using the formula: (X_value_-X_min_)/(X_max_-X_min_).

### Lung function decline is linked to genotypic and phenotypic changes in clinical isolates

Associations between specific bacterial genotypes or phenotypes with patient health during chronic infections lasting years have been uncertain (26,43,44). Many chronic infections are polymicrobial, sampled infrequently and unevenly, and rarely aligned with specific patient variables (23,45,46). Thus, correlations between patient health using a systemic measure like spirometry for lung function and the evolutionary changes of colonizing microbes are challenging to measure and have been limited in their predictive power. Nonetheless, we reconsidered these associations using the breadth and depth of this study. We obtained spirometry measures for seven patients, five of which were temporally aligned with the *B. multivorans* samples (S6A-C Fig and S1 Table). Five patients (P0148, P0213, P0280, P0339, and P0431) exhibited a progressive decrease in lung function based on most measures (S14A Table). Specifically, the percentage of predicted FEV_1_ (forced expiratory volume in 1 second), FVC (forced vital capacity), and FEF_25-75_ (forced expiratory flow at 25-75% of the pulmonary volume) decreased by an average of 4.2%, 10.6%, and 5.3% per year in these patients. Observed lung function for patients P0342 and P0426 remained high during available measures, but these did not overlap the complete duration of isolate collection and limited further study (S6A-C Fig). For all patients with lung function data, we evaluated multiple continuous models for best fit based on Akaike information criterion (AIC) (Fig 7 and S6A-C Fig; S14B Table). Temporal changes in lung function were best fit by a linear decline in two patients (P0342 and P0426), but for five patients, lung functions were better explained by models with varying rates of change. For example, for patient P0213, the cubic model was the best fit (AIC = 343.74, R^2^ = 0.83) with an initial increase in lung function followed by a sharp decline (Fig 7A). Similarly, the lung function of patient P0280 (Fig 7B) showed an initial period of slow decline, followed by a sharp decrease.

**Fig 7.**
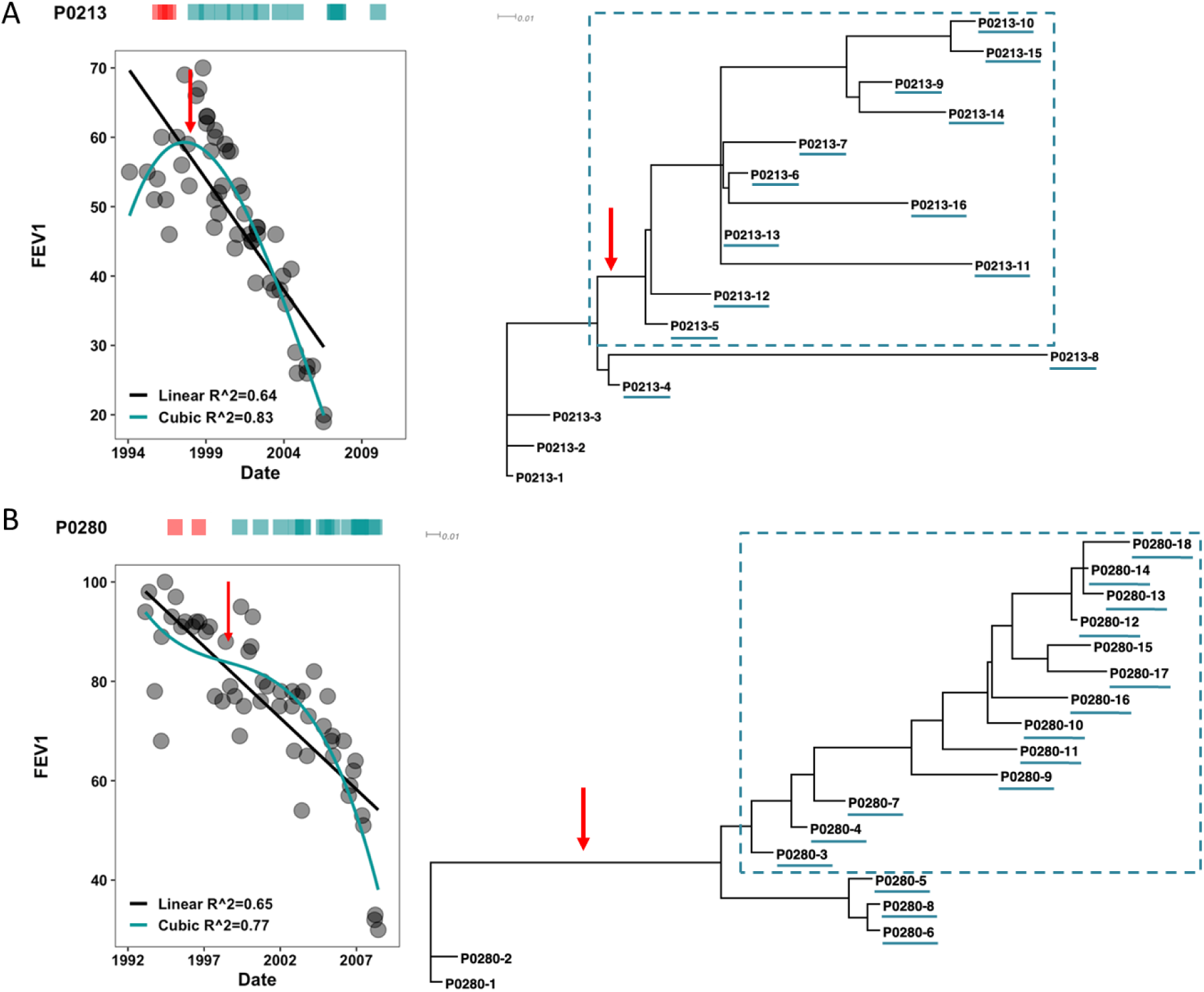
Patient lung function decline is linked to the emergence of dominant lineages within the chronic *B. multivorans* infections. Left: Scatterplots of patient lung function assessed by measuring the forced expiratory volume in 1 second (FEV_1_). The start of lung function decline is denoted by a red arrow. Right: phylogenetic tree modeling SNPs, indels, and large deletions, and the lineages within each longitudinal series (underlined in green) that emerged at the beginning of lung function decline, are shown. Red arrows on the phylogenies denote the estimated timing of the period of patient decline. The dashed rectangle represents the isolates belonging to the dominant clade. The longitudinal series are: (A) P0213, and (B) P0280.

To determine whether periods of accelerated lung function decline coincided with the emergence of new *B. multivorans* lineages, we reexamined phylogenetic trees alongside isolate collection dates (Figs 2-3 and S3A-B Fig). Notably, accelerated periods of lung decline coincided with the time when the *B. multivorans* genetic lineage that would become dominant was first detected. For example, the decline for patient P0213 occurred around the time when isolate P0213-5, an early representative of the dominant lineage, was retrieved (Fig 2A and 7A). In patient P0280, the fitted cubic model describes two distinct periods of worsening lung function: from 1993-1999 annual percentage loss was 2.8% while from 1999-2008 the annual loss was 4.3%. Again, a new dominant lineage was detected at the time of this second period of faster decline (around the time of P0280-3 isolation) (Fig 7B). In both patients, the year in which the dominant lineage was detected (1999) coincided with the onset of greater decline in lung function, an association that remained significant after correcting for multiple tests with empirical *P*<1 x 10^-4^ for FEV_1_. Globally, our data points to the emergence of new dominant *B. multivorans* lineages in chronic respiratory infections coinciding with periods of accelerated lung function decline in infected patients.

We also tested whether the development of pathoadaptive phenotypes shown in Fig 5 correlated with changes in patient lung function (S7 Fig). This analysis revealed strong positive correlation between the three measures of patient lung function and isolate mucoidy, presence of O-antigen, swimming motility, and susceptibility to ciprofloxacin and kanamycin. Thus, in later stages of infections, when patient lung function becomes compromised, isolates tend to exhibit higher resistance to some antimicrobials, lower motility, and the absence of both O-antigen and mucoidy. Overall, these analyzes show that lung function decline likely relates to genotypic and phenotypic changes in the bacterial isolates, in line with previous findings (26).

### Mutations in bacterial genes controlling lipid metabolism and cell wall biogenesis associate with lung function decline

To complement the previous analysis, associations between specific bacterial genotypes and patient lung function were evaluated. Several bacterial mutations were significantly associated with lung function decline (FEV_1_, FVC, FEF_25-75_) (S12 Table). The top five mutated genes correlated with declines in all lung function measurements were *fabF*, *resE*, *plsX, amrR*, and *acrB* encoding a multidrug efflux pump subunit (Fig 8, S12 Table). These findings suggest that phenotypes caused by specific mutations affecting lipid metabolism, antimicrobial resistance, and signal transduction are markers of bacterial adaptation to the CF airway that predict loss of lung function. However, whether and how these *B. multivorans* adaptations contribute to degrading lung function remains uncertain, and it is possible that the decline in airway condition precedes and selects for these bacterial traits.

**Fig 8.**
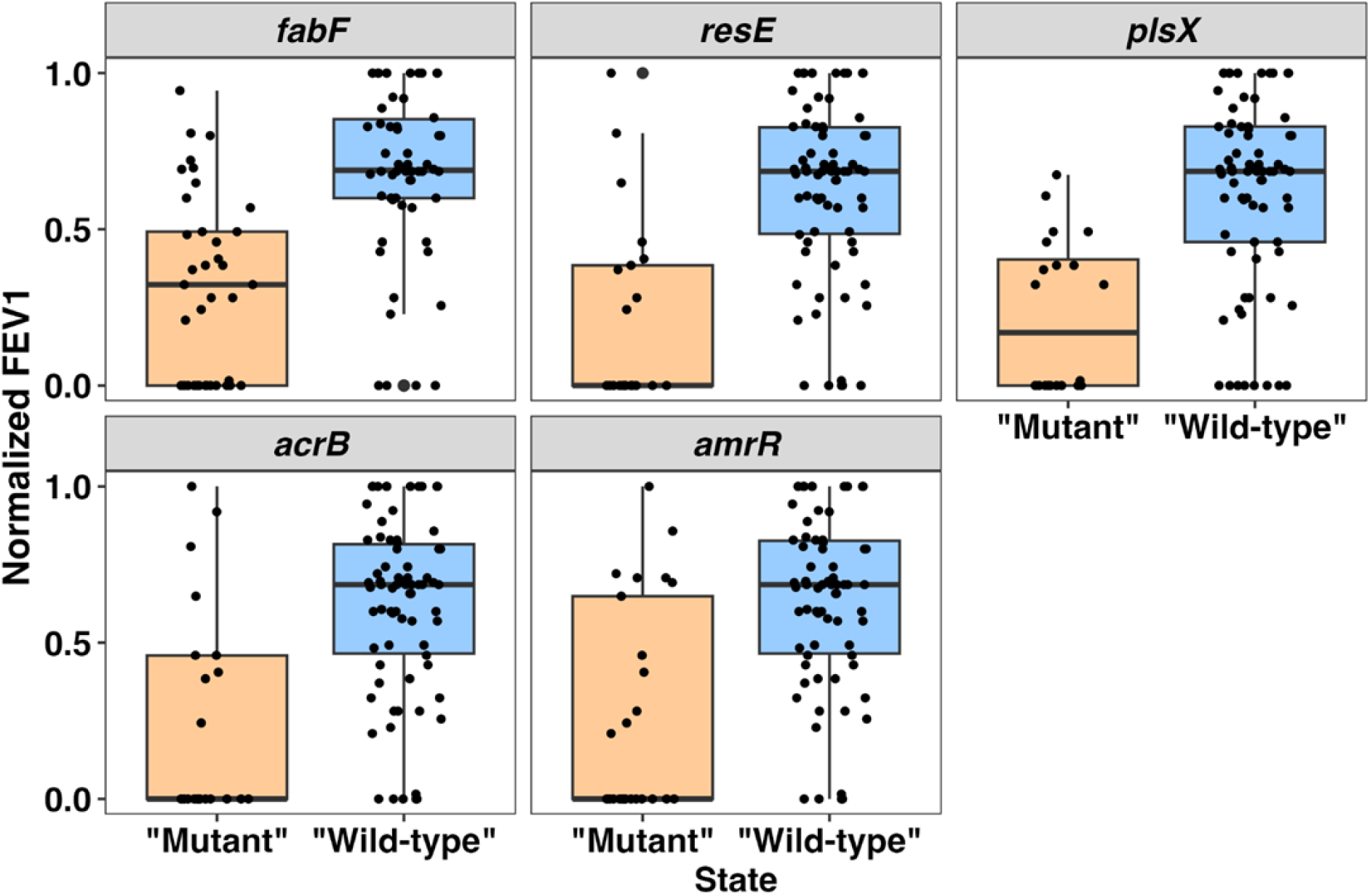
Mutated bacterial genes associated with declines in patient lung function (FEV _1_). Dots represent individual isolates that were categorized as mutant or wild-type depending on the presence or absence of a mutation in the indicated gene. The y-axis depicts normalized FEV_1_ of patients around the time of each isolate collection, determined using the formula: (X_value_-X_min_)/(X_max_-X_min_). Detailed statistical analyses are in S13 Table; each association shown here is supported by *P*<0.0001.

## Discussion

Despite being a monogenic inherited disease, CF is understood by patients and clinicians for its variability in presentation, progression, and symptoms, producing a highly personalized disease history (47). Even the common homozygous F508 CFTR genotype may interact with an unknown number of other variable genes (48), and the atypical microbiome that colonizes their upper and lower airways is typically comprised of unique species and strains (49,50). Therefore, it seems paradoxical that unrelated strains of a diverse environmental species of bacteria would evolve similarly when colonizing different CF patients. Yet, here we show that each *B. multivorans* infection followed broadly similar phylodynamic patterns involving mutations in a small collection of genes producing highly convergent phenotypes. Further, following an initial infection period of 4-5 years when the population underwent limited diversification, one lineage emerged that dominated the subsequent infection history, and this invasion coincided with the accelerated loss of patient lung function in multiple patients. This trend was also seen previously in a single *B. multivorans* CF chronic infection that employed similar methods (26). We suggest these findings are broadly significant for our understanding of both these pathogens and the CF airway environment that may become actionable for diagnostics and treatment strategies.

To begin, why did different *B. multivorans* infections apparently follow similar trajectories? A simple explanation could be sampling bias, because only 6-21 isolates per patient isolated over 7-17 years may have missed important history. Nonetheless, the phylogenetic trees suggest that most major lineages were identified and any undetected variation likely represented rarer lineages that were outcompeted. Otherwise, we would expect to have isolated genotypes outside the dominant lineage in later years (Figs 2-3, S3A-B Fig). Likewise, the absence of singleton or minority genotypes at later points suggests that the early period of genetic diversification disappeared with the sweep of the dominant lineage. These dynamics imply that *B. multivorans* adapted slowly to the CF airway and potentially also to multiple conditions, but eventually one lineage acquired mutations in four or more genes (Figs 4 and 8) that significantly improved fitness throughout the infection and outcompeted other lineages. Among these loci, two central regulators identified previously for their roles in pathogenesis in other *Bc*c, *rpfR* and *fixL,* play major roles, and our analyses point to other genes such as *rstB* and *resE* encoding regulators and the conserved lipoprotein-encoding gene Bmul_1477, discussed below. In summary, these specific mutations appear to enable the rise in frequency and long-term persistence of affected lineages, and thus act as drivers of chronicity and pathogenesis.

The establishment of these dominant lineages appears to have enabled further adaptive evolution, including the convergent evolution of SNPs in many genes (Fig 4) as well as larger deletions. Genome reduction during chronic *Bc*c infections has been reported multiple times and can comprise few to large sets of genes and even entire chromosomes (23,26,51). Nevertheless, large deletions are rarely incorporated in phylogenetic reconstruction due to algorithmic complexity. Including these events in our models showed that large deletions arise predominantly after the formation of dominant clades. This suggests early adaptations to the CF lung are produced by SNPs and small indels, whereas larger deletions tend to occur in better adapted strains, potentially owing to population bottlenecks and reduced selection in established infections. Indeed, most deletions involved hypothetical proteins associated with mobile elements, and deletions affecting virulence factors, e.g. in regulators or in O-antigen biosynthesis, tended to occur in single isolates or in lineages that had already become deficient in O-antigen production. Both trends are consistent with effects of drift more than selection. Our results also quantitatively agree with studies of chronic *P. aeruginosa* infections in CF that experience deletions of an average of 88 genes (52). This dynamic of genome erosion is consistent with declining purifying selection in bottlenecked pathogen populations or new animal symbionts (12), although some gene losses may produce adaptations such as the evasion of the host immune response by losing virulence factors (53).

Integrating our study with prior studies of *Bc*c infections builds understanding of how these bacteria adapt to the CF airway and what host conditions exert selection. Using the strict criterion of orthologous genes mutated in at least two patients infected by all three of the best studied *Bc*c species –*B. multivorans* (26–28,54), *B. cenocepacia* (51,54), and *B. dolosa* (21,22) –we identify nine genes as clear evidence of repeatable evolution in the CF airway (S15 Table). When we broaden this analysis to pathways, the most commonly affected were lipopolysaccharide biosynthesis (e.g., *wbi*, *wbbL*, *rmd*, *lptB*), antibiotic resistance (e.g., *rpoB*, *rpoA, mpl, gyrA*) and siderophore biosynthesis or transport (e.g., *orbI*, *orbJ*, *pvdA*). Mutated genes shared between this study and those of *B. dolosa* adds convergent changes in lipid metabolism (e.g., *fabF*, *fabG*) and global regulation (*vnfA* and *rstB*). Likewise, mutations shared with longitudinal samples of *B. cenocepacia* identifies additional global regulators (*rpfR*, *resE*, and *relA*/*spoT*). Therefore, despite considerable methodological differences, with some sequencing clones of epidemic strains and others performing deep sequencing directly from sputum samples, *Bc*c strains adapt to the CF airway by a common set of pathways and phenotypes. Collectively, this work depicts a scenario where *Bc*c must acquire mutations in five or more specific pathways to get a ‘winning hand’ that establishes chronic infection and ultimately degrades the airway. Fewer mutations are insufficient, like having only a pair in a game of poker when you need a full house. Further, there appears to be an approximate order in these evolved *Bc*c traits, which begin with mutated global regulators, then altered lipid metabolism, immune evasion, antibiotic resistance, stringent response, iron acquisition, and survival under oxygen limitation, the latter of which suggest survival either intracellularly or in nearly anoxic pockets. Monitoring when these core adaptations occur in the establishment and succession of *Bc*c chronic infections could be clinically valuable for diagnostics or management.

Returning to the poker analogy, the global regulators that appear to be essential components of a winning hand are *rpfR, fixL,* and *rstB*. In five patient series, mutations in *rpfR* preceded the emergence of the lineage that became dominant in each infection and also prior to mutations in *fixL* and *rstB* genes, both encoding sensor kinases. RpfR is a global regulator for *Bc*c and related bacteria because it integrates sensing of the external environment through quorum sensing with the motile-sessile switch mediated by cyclic-di-GMP (39). RpfR regulates many virulence factors, such as the production of the polysaccharide Bep, biofilm formation, motility, and others (39,40,55). Some of these phenotypes are also influenced by FixL/FixJ (36) and RstA/RstB, which govern responses to hypoxia and stress responses (56). Our data suggest that RpfR sits atop a regulatory cascade involving these two signal transduction systems, but this requires further study.

Identifying microbial traits as biomarkers of worsening chronic infection is a broadly held goal of modern clinical microbiology. Culture-based bacterial phenotypes remain a more accessible indicator, but the variability of bacterial isolates from sputum or lavage samples has confounded predictors (23,57). We also observed phenotypic diversity among isolates within and between patients, but most phenotypes changed in a common direction over time. Early isolates were characterized by higher growth rate, presence of O-antigen, mucoid colonies, higher motility, higher acute virulence when tested in an animal model, and increased antibiotic susceptibility, while evolved isolates tend to have the opposite phenotypes. This does not mean early phenotypes cannot reoccur later in the infection. For example, in patient P0426, some later isolates of clades C5 and C6 likely recovered O-antigen, as seen in *B. dolosa* (58), while displaying other chronic-like traits such as slow growth and antibiotic resistance. Evolution in the heterogeneous CF environment both selects and maintains phenotypic diversity that contributes to persistence in the face of stress. Nonetheless, the convergent phenotypes and genotypes observed here may serve as markers of progression towards chronicity for new patients with *Bc*c infections (59,60).

Our study has certain limitations given its retrospective nature. First, we lack consistent data about other co-infecting species, which renders it impossible to determine if the dominant clade of *B. multivorans* is responsible for, or a result of, reduced lung microbial diversity. Nonetheless, many studies have reported a correlation between a decline in lung function and a loss of microbial diversity (45,61). Second, at most time points, only a single isolate is available. Isolates therefore reflect the cultured fraction of the sputum community where media choice and colony appearance may have influenced lineage recovery. However, because consistent culturing and sequencing methods were applied across seven independent chronic infections, the repeated observation of early diversification followed by later consolidation of a dominant clade supports a robust qualitative phylodynamic pattern among cultured isolates. Third, we did not employ recombination-aware phylogenetic methods that could alter our inference of a small fraction of within-host parallel evolution. Nonetheless, the consistency of tree topologies between parsimony and maximum-likelihood inference, as well as across different character sets (SNPs-only, SNPs+indels, and SNPs+indels+large deletions (S8 Fig)), strengthens confidence in the robustness of the identified dominant clade and the inferred within-host evolutionary dynamics. Lastly, we could not conduct a true genome-wide association study due to insufficient sampling and the clonal population structure of bacterial infection. However, our genotype-phenotype correlations revealed associations that we could test experimentally that validates our approach (S13 Table). Future studies can benefit from more sampling, functional validation, and phylogenetic context correction to strengthen mechanistic understanding of genetic determinants of pathogenic phenotypes.

The convergent evolution we observe across independent *B. multivorans* CF infections –involving a conserved set of approximately five key genes repeatedly mutated in a predictable sequence –reveals that the CF airway environment exerts powerful selective pressures that constrain *Bc*c adaptation to a limited number of solutions. This stereotyped adaptive trajectory, combined with the temporal dynamics of dominant lineage emergence coinciding with accelerated lung function decline, provides a novel framework for monitoring and managing chronic *Bc*c infections. Specifically, mutations in driver genes (*rpfR*, *fixL*, *plsX*, *amrR*) could serve as early diagnostics of infection establishment and chronicity, enabling stratified clinical intervention before substantial airway damage occurs. Furthermore, these convergent pathways represent rational therapeutic targets that could prevent the transition from acute to chronic infection or restore antibiotic susceptibility in established infections. Comparative analyses across the *Bc*c species demonstrates that these conclusions extend broadly across *Bc*c pathogens, suggesting that therapeutic strategies targeting these core adaptive mechanisms could have broad applicability beyond *B. multivorans* and potentially inform management of other chronic CF infections.

## Conclusion

Our findings reveal a highly convergent evolutionary route by which *B. multivorans* adapts to the cystic fibrosis airway, with recurrent changes in global regulation and linked phenotypes that track the emergence of dominant lineages and worsening lung function. These results are broadly relevant to other chronic infections by environmental opportunists, show that pathogen population dynamics can be clinically relevant markers of chronic infection progression, and identify conserved adaptive pathways as promising targets for earlier diagnosis and intervention.

## Materials and methods

### Bacterial strains and growth conditions

Clinical isolates listed in S1 Table were obtained from the CBCCRRR. Stocks two passages away from the originals were prepared, and all subsequent manipulations used a single passage from these stocks. *Burkholderia* isolates were routinely grown at 37°C in LB agar, and liquid cultures were grown in LB or SM medium (62).

### Ethics statement and patient data collection

The bacterial samples and lung function dataused in the study were approved by the University of British Columbia Research Ethics Boards with reference H07-01396 (43). The committee determined that informed consent was not required because the study used pre-existing clinical isolates and routinely collected data that were de-identified prior to researcher access. No directly identifying participant information was accessed or analyzed.

### DNA extraction and genome sequencing

Genomic DNA for Illumina sequencing was extracted from an overnight culture of each *B. multivorans* isolate using the DNeasy blood and tissue kit (Qiagen, Germany) following the manufacturer’s instructions for Gram-negative bacteria. Multiplexed sequencing libraries made with the NEXTera XT DNA sample prep kit (Illumina) and sequencing to obtain paired-end reads to a minimum read depth of 50x were performed as a service at the Genomics Unit of Instituto Gulbenkian de Ciência, Portugal. Genomic DNA for PacBio sequencing was extracted using the Qiagen genomic-tip 20/G kit following the manufacturer’s instructions. A 20-kb SMRTbell library was prepared, and sequencing was performed using SMRT cells with P4-C2 chemistry at the Icahn School of Medicine at Mount Sinai, USA.

### *De novo* genome assembly and annotation

Illumina raw paired-end reads from *B. multivorans* isolates were filtered with the trimming tool sickle (version 1.33) with a quality score of 30 (63). *De novo* assembly was carried out with SPAdes (version 3.12.0) with k-mer lengths of 21, 33, 55, 77, and in the “only assembler” mode (64).

For PacBio-sequenced isolates P0213-1, P0280-1, and P0426-1, the reads were assembled with the HGAP assembler using the SMRT Analysis Suite (Pacific Biosciences) (65). These three assemblies were aligned to a reference-*B. multivorans* ATCC 17616 - to reorder contigs using Medusa (v1.6). Then, Illumina reads of the same isolates were trimmed with Trimmomatic and aligned to medusa-assemblies with BWA (v0.7.15). The mapped reads were used to predict and correct misassemblies with Pilon (v1.19). All assembled genomes were annotated with the rapid prokaryotic genome annotation pipeline, Prokka, v1.12 (66). In S17 Table can be found the details of assembly statistics of reads and the number of Prokka-predicted genes.

### MLST analysis

Estimation of the sequence type (ST) of each isolate by multi-locus sequence typing was done using the Short Read Sequence Typing for Bacterial Pathogens (SRST2) v0.2.0 tool (67). This program uses filtered Illumina sequence data and the MLST database for the *Burkholderia cepacia* complex (https://pubmlst.org/organisms/burkholderia-cepacia-complex) and reports on the ST identified.

### Population structure of *B. multivorans*

The core genome alignment of *B. cenocepacia* K56-2 and 123 *B. multivorans* reference strains, and the first isolates of each *B. multivorans* longitudinal series under study (S3 Table) was used to extract the core genome SNPs by using the SNP sites program v2.0.2 from the Sanger Institute (https://github.com/sanger-pathogens/snp-sites). These SNPs were used as input for FastTree (http://www.microbesonline.org/fasttree/) v2.1.11 (68) to infer approximately-maximum-likelihood phylogenetic tree that was visualized with FigTree v1.4.4..

### Mobilome analysis

Prediction of genomic island content was done with the IslandCompare tool (69). The Phaster tool was used for detecting prophages regions (70,71) and the mobileOG-db tool for detection of various mobile genetic elements (72). Antibiotic resistance genes were searched against the Comprehensive Antibiotic Resistance Database (CARD) (73). Graphics overlapping mobile elements with deleted regions in each isolate genome were obtained using Proksee software (74).

### Pan-genome analysis

Pan-genome analyzes for all isolates of each longitudinal series were performed using the rapid large-scale prokaryote pan-genome analysis pipeline, Roary **version 3.11.2,** from the Sanger Institute (75). Depending on the analyzes, several NCBI reference genomes were included. The input files were the GFF3 files from the Prokka-annotated assemblies of our isolate’s genomes, and the GFF created from the GenBank format for the reference genomes included in S17A Table. To create core-genome alignments, Roary first clustered coding regions using CD-HIT (76), followed by an all-against-all comparison between the coding regions using a minimum cutoff of 75% BLASTP identity, ignoring truncated and un-annotated genes, and genes with fewer than 120 bases. By joining the BLAST results with MCL clustering (77), Roary identified the homologous groups of genes, with a codon-aware multiple sequence alignment generated by PRANK (78) for both the core genome and the accessory genome.

### Detection of SNP and indel mutations

Filtered paired-end datasets were mapped to the reference genome from the first isolate of each longitudinal series using BWA-MEM v0.7.10 (79) and NovoAlign v3.02.13 (Novocraft technologies). Single-nucleotide variants were called and filtered with SAMtools v0.1.18 (80), and predicted SNPs and indels were manually inspected for coverage, allele frequency, and strand support in Geneious v6.1.8 (81).

### Temporal-signal assessment by root-to-tip regression

To assess temporal signal, maximum-likelihood trees were analyzed in TempEst v1.5.3 (82) using the earliest isolate as the root and reference genome. Root-to-tip regression of genetic distance against sampling date was evaluated using the residual mean squared criterion, and slope, R², correlation coefficient, and TMRCA were recorded. The analysis was restricted to patient series without hypermutators, except for P0313-8.

### Patient series phylogenetic analysis

To infer the phylogeny of a patient series, SNPs, indel mutations, and large deletions were concatenated separately for each isolate to create a GenomeDiff file. The command COMPARE included in the gdtools utility program from breseq (83) (https://barricklab.org/twiki/pub/Lab/ToolsBacterialGenomeResequencing/documentation/gd_usage.html) merged all GenomeDiff files creating a Phylip file as an output. Then, the PHYLogeny Interface Package (PHYLIP) v3.695 (84) program dnapars was used to create a maximum parsimony phylogenetic tree file. To visualize the outree file (in Newick format), tool TreeDyn 198.3 (http://www.phylogeny.fr/one_task.cgi?task_type=treedyn) was used. Phylogenetic reconstruction initially used a parsimony-based approach to capture genome-wide variation across longitudinal isolates, integrating SNPs, indels, and large structural changes with minimal assumptions. This framework enabled identification of broad evolutionary relationships and diversification patterns while incorporating multiple classes of variation difficult to model explicitly. Maximum-likelihood (ML) phylogenetic inference was performed to assess the robustness of inferred topologies, using explicit nucleotide substitution models and bootstrap support. Concordance between ML and parsimony trees, as well as across datasets (SNPs-only, SNPs+indels, SNPs+indels+large deletions), supports the stability of the inferred relationships.

### Quantification of clade dominance and temporal structure

For quantitative bounds on dominance and on the likelihood of observing the reported patterns under sparse sampling, we re-analyzed per-time point data where ≥2 isolates were collected to estimate clade frequencies and exact 95% Clopper-Pearson confidence intervals. Then, binomial sensitivity analyses for time points with n = 1-4 to quantify the probability of missing minority clades at plausible underlying frequencies (q = 0.1-0.3) were performed. Additionally, computed lineage ‘survival’ curves (Kaplan-Meier) using the interval between first and last detection to summarize the persistence of early clades was conducted. Finally, a permutation test (10,000 permutations per patient) that randomized clade labels across the observed sampling dates to estimate how often exclusive late recovery of a single clade would occur by chance under the empirical sampling scheme was run (S8 Table).

### Gene-level recurrence analysis in non-hypermutator series

Series-level neutral substitution rates were estimated by combining *in vitro* doubling times in SCFM medium (S1 Table) for each non-hypermutator longitudinal series with a baseline mutation rate of 1.33×10^-10^ substitutions/base/generation (38), converting each isolate’s doubling time to generations/year and then to substitutions/base/year. The series-specific rates were summarized by their medians, yielding five values between 5.7×10^-7^ and 1.12×10^-6^ substitutions/base/year. We averaged these to obtain a mean neutral rate of 7.58×10^-7^ substitutions/base/series, and for each of 29 genes (mutated in ≥2 of 5 non-hypermutator series) used gene length and this neutral rate to compute the per-series hit probability *p_i_* = 1-*e^-μL^*, the expected number of hit series *E*[*X_i_*] = 5*p_i_*, and exact one-sided binomial *P*-values *P*(*X_i_* ≥ *k_i_*), with Benjamini-Hochberg FDR applied across genes. All gene-level statistics are provided in S10J Table.

### Genotype-phenotype associations

To link phenotypic variation with adaptive mutations, two matrices were generated: a normalized phenotypic matrix (xvalue-xmin)/(xmax-xmin) and a binary genotype matrix (“1” wild-type, “0” mutant). The “mutant” gene is considered being any mutation, including those affecting intergenic regions (SNPs and indels) or coding sequences, which can be synonymous, nonsynonymous, indels, or large deletions. The “wild-type” alleles were defined relative to the reference genome. Roary gene families ensured non-redundant homolog grouping. Spearman’s rank correlation was then applied to identify associations between genotype and phenotype.

### Patient-level permutation analysis of lineage detection and accelerated decline

Patient-level permutation tests were used to evaluate whether detection of the dominant lineage coincided with accelerated lung-function decline more often than expected by chance. For that, longitudinal lung-function values were randomly reassigned among collection times 10,000 times, and empirical *P*-values were adjusted for the outcomes using the Benjamini-Hochberg procedure.

### Determination of growth rates and doubling time

Cultures were grown in SCFM (85) at 37 °C, 200 rpm orbital agitation, and OD_640nm_ was measured for 24 hours. The doubling time was calculated from the growth rate of the exponential growth phase. Three independent experiments with two replicates each were performed.

### Mucoid phenotype determination

Mucoid colony morphology was assessed in yeast extract-mannitol (YEM) agar medium (0.5 g/l yeast extract; 4 g/l D-mannitol; 2% agar) by incubating inoculated plates at 37°C for 48 hours. Mucoid level estimation followed the procedure of Zlosnik and co-authors (29).

### Analysis of the lipopolysaccharide

Lipopolysaccharide extraction and purification was performed by the method of Marolda and co-authors (86) from 1.5 ml of *B. multivorans* cell suspensions with OD_600nm_ of 2.0. The analysis of the LPS O-antigen samples was carried out by electrophoresis in 16% polyacrylamide gels using Tricine-SDS and visualized by silver staining. The analysis of the LPS O-antigen samples was carried out by electrophoresis in 16% polyacrylamide gels using Tricine-SDS and visualized by silver staining.

### Antimicrobial susceptibility

Antimicrobial susceptibility was based on the agar disc diffusion method (87) against piperacillin (75 μg) plus tazobactam (10 μg), aztreonam (30 μg), ciprofloxacin (5 μg), and kanamycin (30 μg). The discs, obtained from Becton Dickinson, were applied onto the surface of Mueller-Hinton agar plates (Difco Laboratories) that had been previously inoculated with 100-μl aliquots prepared from cultures grown overnight in LB, at 37°C, with agitation, and diluted to a standardized culture OD_640nm_ of 0.1. Diameters of growth inhibition were measured after 24 h of incubation at 37°C. At least three independent experiments, each containing four technical replicates, were performed.

### Motility assays

For motility estimation, 5 μl of overnight LB bacterial cultures were inoculated onto the agar surface of swimming and swarming plates, and incubated statically at 37°C for 24 and 48 hours, respectively, followed by colony diameter determination. Swimming plates were prepared with 1% (wt/vol) tryptone, 0.5% (wt/vol) NaCl, 0.3% (wt/vol) noble agar (Difco) while swarming plates were prepared in Broomfield medium (0.04% (w/v) tryptone, 0.01% yeast extract (w/v), 0.0067% (w/v) CaCl_2_) with 0.6% (w/v) of Bacto agar. Three independent experiments, each containing four technical replicates, were performed.

### Biofilm formation on abiotic surfaces

Biofilm assays were performed by the method of Ferreira and co-authors (88). Overnight liquid cultures of the different *B. multivorans* strains were diluted to a standardized culture OD_640nm_ of 0.1 in LB and 200 μl of these cell suspensions were inoculated into the wells of a 96-well polystyrene microtiter plate. Plates were incubated at 37°C for 48 h without agitation. The biofilm was stained with crystal violet solution, followed by dye solubilization with ethanol and measurement of the solutiońs at OD_590nm_ using a microplate reader. Results are mean values for at least five repeats from three independent experiments.

### Host cell attachment

*B. multivorans* isolates were analyzed for adhesion to the bronchial epithelial cell line CFBE41o-derived from a patient homozygous for the cystic fibrosis transmembrane conductance regulator F508del mutation (89). Bacterial strains were grown overnight in LB medium, after which 200 µl of those cultures were grown in LB for 4 hours and then used to infect epithelial cells at a multiplicity of infection (MOI) of 10 (10 bacterial cells to 1 epithelial cell). The experimental procedure was according to (90). Duplicates of each strain were performed per assay, and the results presented were obtained from three independent experiments. Results are shown as the percentage of adhesion, which was calculated as the number of CFU recovered divided by the number of CFU applied to the epithelial cells multiplied by 100.

### Virulence determination in *Galleria mellonella*

Killing assays were performed as previously described (91). Larvae were injected with 1×10^6^ CFU diluted in 10 mM MgSO_4_ with 1.2 mg/ml ampicillin, and the survival rate was evaluated at 24-, 48- and 72-hours post-infection. As a negative control, 10 mM MgSO_4_ with 1.2 mg/ml ampicillin was used. This assay was repeated four times with ten larvae each.

### Statistical analysis

All quantitative data were obtained from at least three independent assays with at least two biological replicates. Differences were assessed by one-way ANOVA with Dunnett’s or Tukey’s multiple comparisons test, and by the Mantel-Cox test, using GraphPad Prism v.7.02 (GraphPad Software, USA) and R (92). *P<*0.05 was considered statistically significant.

### Data and materials availability

The DNA sequence reads for assemblies of the several *B. multivorans* genomes have been submitted to the NCBI BioProject under accession number PRJNA542030 and can be assessed from the Sequence Read Archives (SRA) with accession numbers SRR9046870-SRR9046975. Additional data are available in the main text or the supplementary materials.

## Acknowledgments

We thank Dr. David P. Speert and the Canadian *Burkholderia cepacia* Complex Research and Referral Repository (CBCCRRR) from the University of British Columbia, Vancouver, Canada, for the *Burkholderia multivorans* isolates. We would also like to thank Dr. Pedro Santos from the University of Minho for the helpful discussions on genomic data analysis.

## Supporting information

**S1 Fig. Core genome SNP phylogeny across selected *B. multivorans* isolates.** Unrooted phylogeny was built using FastTree from the 3035 orthologous clusters of protein-coding genes from the 9 isolates (P0148-1, P0213-1; P0280-1; P0339-1; P0342-1; P0426-1; P0431-1a; P0431-2b; and P0686-2b, shown in red) and 123 NCBI reference strains (S3 Table). Isolates P0431-1a, P0431-2b, and P0686-2b are represented in the figure by the names P0431-1, P0431-2, and P0686-2, respectively. The genome of *B. cenocepacia* K56-2 was used as an outgroup. Clade 1 strains are shown in blue and clade 2 strains in black.

**S2 Fig. Within-host SNP accumulation and root-to-tip analyses across longitudinal *Burkholderia* series showed a temporal signal.** (A) Number of SNPs distinguishing each isolate from the first of the indicated series over time. Mutator isolates with mutations in *mutS* or *mutL* genes (series P0213 and P0431a, shown in red) were excluded from the estimation of mutation rates. Green dots also represent isolates excluded from the analysis because of being less evolved than earlier isolates (series P0213) or from being to a significantly different lineage (series P0342). In P0342 series, isolates with mutations in the *mutT* gene (shown in orange) were not excluded from the analysis because they share mutations in other DNA repair genes in common with the previous isolates (shown in blue). The x and y axis legend are the same for each chart. (B) Temporal signal of the phylogeny inferred from maximum-likelihood trees using TempEst v1.5.3. The phylogeny was rooted using the earliest chronological isolate, which also served as the reference genome for variant calling and within-host evolutionary analyses. Root-to-tip regression plotted genetic distance from the root against sampling date. The regression was evaluated using the Residual Mean Squared criterion, and the slope, R², correlation coefficient, and estimated time to the most recent common ancestor (TMRCA) were recorded to assess temporal signal. This analysis was performed only for patient series without hypermutator isolates, except P0313-8, because accelerated mutation rates in hypermutators violate the approximately constant-rate assumption of root-to-tip regression.

**S3A Fig. Genomic phylogeny reveals the long-term coexistence of diverse clades.** (**A, B**) Phylogenetic trees of the isolates of P0148 and P0426 longitudinal series, respectively, modeling both SNPs, indels and large structural variations in the genome. The main branches are represented by capital letters, as defined in S10 Table. Symbols indicate the phylogenetic position of putative driver mutations that occurred in the genes shown at bottom right. (**C, D**) Temporal distribution of isolates sampling and respective clades (C_1-3_ in P0148 and C_1-6_ in P0426 series) shows that clades coexisted. Symbol color indicates clade membership, with line color indicating major clade and fill color indicating sub-clade.

**S3B Fig. Genomic phylogeny reveals the long-term coexistence of diverse clades.** (**A, B, C**) Phylogenetic trees of the isolates of P0431, P0342 and P0339 longitudinal series, respectively, modeling both SNPs, indels and large structural variations in the genome. The main branches are represented by capital letters, as defined in S10 Table. Symbols indicate the phylogenetic position of putative driver mutations that occurred in the genes shown at bottom right. (**D, E, F**) Temporal distribution of isolates sampling and respective clades (C_1-5_ in P0431, C_1-2_ in P0342, and C_1-2_ in P0339 series) shows that clades coexisted. Symbol color indicates clade membership, with line color indicating major clade and fill color indicating sub-clade.

**S4A Fig. Genome reduction in *B. multivorans* longitudinal series is enriched in mobile genetic elements.** Comparison of the serial isolates’ genomes against the reference genome of the first isolate depicting the major deletions merged with the location of the mobile genetic elements in P0148, P0280, and P0426 longitudinal series. Starting from the outermost ring: (Ring 1) genomic islands; (Ring 2) prophages; (Ring 3) mobile elements. Graphics obtained using Proksee software.

**S4B Fig. Genome reduction in *B. multivorans* longitudinal series is enriched in mobile genetic elements.** Comparison of the serial isolates’ genomes against the reference genome of the first isolate depicting the major deletions merged with the location of the mobile genetic elements in P0213 and P0339 longitudinal series. Starting from the outermost ring: (Ring 1) genomic islands; (Ring 2) prophages; (Ring 3) mobile elements. Graphics obtained using Proksee software.

**S4C Fig. Genome reduction in *B. multivorans* longitudinal series is enriched in mobile genetic elements.** Comparison of the serial isolates’ genomes against the reference genome of the first isolate depicting the major deletions merged with the location of the mobile genetic elements in P0342, P0431a and P0431b and longitudinal series. Starting from the outermost ring: (Ring 1) genomic islands; (Ring 2) prophages; (Ring 3) mobile elements. Graphics obtained using Proksee software.

**S5 Fig. Phenotypic variation along the series of *B. multivorans* clinical isolates recovered from seven CF patients.** A *P*-value < 0.1 was considered significant.

**S6A Fig. Scatterplots of patient lung function assessed by measuring the forced expiratory volume in 1 second (FEV_1_).** Stated above each chart (green squares) is the time point of each *B. multivorans* collected isolate from each longitudinal series. The linear model and the model that best fits the data (with the lowest AIC value, S14B Table) were plotted on each graph.

**S6B Fig. Scatterplots of patient lung function assessed by measuring the forced vital capacity (FVC).** Stated above each chart (green squares) is the time point of each *B. multivorans* collected isolate from each longitudinal series. The linear model and the model that best fits the data (with the lowest AIC value, S14B Table) were plotted on each graph.

**S6C Fig. Scatterplots of patient lung function assessed by measuring the forced expiratory flow at 25-75% of the pulmonary volume (FEF_25-75_).** Stated above each chart (green squares) is the time point of each *B. multivorans* collected isolate from each longitudinal series. The linear model and the model that best fits the data (with the lowest AIC value, S14B Table) were plotted on each graph.

**S7 Fig. Correlation matrix of *B. multivorans* isolates phenotypes and patient lung function**. A pairwise Spearman rank correlation was used, with the blue-to-green gradients showing a negative correlation and the green-to-yellow gradients a positive correlation. Correlation values (r_s_) and corresponding *p*-value: ± 0.401-1.0 (*P*-value < 0.0001); ± 0.339-0.401 (*P*-value < 0.001); ± 0.266-0.339 (*P*-value < 0.01); and ± 0.202-0.266 (*P*-value < 0.05).

**S8A Fig. Phylogenetic trees of P0148-1 longitudinal series inferred using a maximum-likelihood framework.** These were implemented in IQ-TREE, with substitution model selection performed by ModelFinder and branch support evaluated using 1,000 ultrafast bootstrap replicates. Each figure shows three phylogenetic trees performed with all mutations (SNPs+indels+large deletions (>2kbp)), only SNPs and SNPs+indels, respectively.

**S8B Fig. Phylogenetic trees of P0213-1 longitudinal series inferred using a maximum-likelihood framework.** These were implemented in IQ-TREE, with substitution model selection performed by ModelFinder and branch support evaluated using 1,000 ultrafast bootstrap replicates. Each figure shows three phylogenetic trees performed with all mutations (SNPs+indels+large deletions (>2kbp)), only SNPs and SNPs+indels, respectively.

**S8C Fig. Phylogenetic trees of P0280-1 longitudinal series inferred using a maximum-likelihood framework.** These were implemented in IQ-TREE, with substitution model selection performed by ModelFinder and branch support evaluated using 1,000 ultrafast bootstrap replicates. Each figure shows three phylogenetic trees performed with all mutations (SNPs+indels+large deletions (>2kbp)), only SNPs and SNPs+indels, respectively.

**S8D Fig. Phylogenetic trees of P0339-1 longitudinal series inferred using a maximum-likelihood framework.** These were implemented in IQ-TREE, with substitution model selection performed by ModelFinder and branch support evaluated using 1,000 ultrafast bootstrap replicates. Each figure shows three phylogenetic trees performed with all mutations (SNPs+indels+large deletions (>2kbp)), only SNPs and SNPs+indels, respectively.

**S8E Fig. Phylogenetic trees of P0342-1 longitudinal series inferred using a maximum-likelihood framework.** These were implemented in IQ-TREE, with substitution model selection performed by ModelFinder and branch support evaluated using 1,000 ultrafast bootstrap replicates. Each figure shows three phylogenetic trees performed with all mutations (SNPs+indels+large deletions (>2kbp)), only SNPs and SNPs+indels, respectively.

**S8F Fig. Phylogenetic trees of P0426-1 longitudinal series inferred using a maximum-likelihood framework.** These were implemented in IQ-TREE, with substitution model selection performed by ModelFinder and branch support evaluated using 1,000 ultrafast bootstrap replicates. Each figure shows three phylogenetic trees performed with all mutations (SNPs+indels+large deletions (>2kbp)), only SNPs and SNPs+indels, respectively.

**S8G Fig. Phylogenetic trees of P0431-1 longitudinal series inferred using a maximum-likelihood framework.** These were implemented in IQ-TREE, with substitution model selection performed by ModelFinder and branch support evaluated using 1,000 ultrafast bootstrap replicates. Each figure shows three phylogenetic trees performed with all mutations (SNPs+indels+large deletions (>2kbp)), only SNPs and SNPs+indels, respectively.

**S1 Table. Patients’ clinical data and phenotypes of *B. multivorans* isolates collected from the 8 cystic fibrosis patients, including date of isolation and RAPD type, if available.**

**S2 Table. Multi-locus sequence type (MLST) of each clinical isolate under study.**

**S3 Table. List of strains with corresponding genomic data used for the phylogenetic tree construction based on the core genome SNPs.**

**S4 Table. Genomic content of isolates from all longitudinal series mobile elements, prophage regions and genomic islands.**

**S5 Table. List of strainś genomes used to identify unique genes in each of them.**

**S6 Table. List of mutations found in all isolates when compared to the first of the longitudinal series.**

**S7 Table. Number of large deletions (> 2 kb) in the longitudinal series collected from CF patients.**

**S8 Table. Temporal statistics describing Burkholderia clonal dynamics across chronically infected patients (n = 93 isolates).**

**S9 Table. Gene presence_absence as determined by Roary (75% BLASTP identity, merged orthologous clusters) for the longitudinal series under study.**

**S10 Table. Enrichment of nonsynonymous mutations and identification of parallel mutations.**

**S11 Table. Patient lung function and phenotypic data normalized mean values obtained using the formula: (xvalue-xmin)/(xmax-xmin) and binary genotype matrix defining each gene as “wild-type”** (**1**) **or “mutant”** (**0**).

**S12 Table. Spearman rank correlations between phenotypes and mutations.**

**S13 Table. Associations between specific mutated genes and clinically relevant phenotypes meeting stringent statistical significance.**

**S14 Table. Linear regression of patient lung function over time and statistic analysis of continuous models used to fit data.**

**S15 Table. Genes mutated in the longitudinally sampled *B. multivorans* infections under study and their discovery in other longitudinal CF infections reported in the literature.**

**S16 Table. Pangenome analysis across longitudinal series of *B. multivorans*.**

**S17 Table. De novo assembly statistics of reads generated by Illumina or PacBio sequencing.**

